# Neural and subjective effects of inhaled DMT in natural settings

**DOI:** 10.1101/2020.08.19.258145

**Authors:** Carla Pallavicini, Federico Cavanna, Federico Zamberlan, Laura Alethia de la Fuente, Yonatan Sanz Perl, Mauricio Arias, Celeste Romero, Robin Carhart-Harris, Christopher Timmermann, Enzo Tagliazucchi

## Abstract

**Background:** N,N-Dimethyltryptamine (DMT) is a short acting psychedelic tryptamine found naturally in many plants and animals. Few studies to date addressed the neural and psychological effects of DMT alone, either administered intravenously or inhaled in freebase form, and none conducted in natural settings.

**Aims:** Our primary aim was to study the acute effects of inhaled DMT in natural settings, focusing on questions tuned to the advantages of conducting field research, including the effects of contextual factors (i.e. “set” and “setting”), the possibility of studying a comparatively large number of subjects, and the relaxed mental state of participants consuming DMT in familiar and comfortable settings.

**Methods:** We combined state-of-the-art wireless electroencephalography (EEG) with psychometric questionnaires to study the neural and subjective effects of naturalistic DMT use in 35 healthy and experienced participants.

**Results:** We observed that DMT significantly decreased the power of alpha (8-12 Hz) oscillations throughout all scalp locations, while simultaneously increasing power of delta (1-4 Hz) and gamma (30-40 Hz) oscillations. Gamma power increases correlated with subjective reports indicative of mystical-type experiences. DMT also increased/decreased global synchrony and metastability in the gamma/alpha band, and resulted in widespread increases in signal complexity.

**Conclusions:** Our results are consistent with previous studies of psychedelic action in the human brain, while at the same time suggesting potential EEG markers of mystical-type experiences in natural settings, thus highlighting the importance of investigating these compounds in the contexts where they are naturally consumed.

## Introduction

Recent years have seen a major renaissance in the neuroscientific and clinical study of serotonergic psychedelics (Sessa, 2018). Even though compounds such as psilocybin, lysergic acid diethylamide (LSD), and N,N-dimethyltryptamine (DMT) were studied during the ‘50s and ‘60s, the modern study of psychedelics has been revitalized by improved experimental designs and ethical standards, as well as by the availability of more sophisticated non-invasive neuroimaging techniques (Lee and Shlain, 1992; Carhart-Harris and Goodwin, 2017). The most robust and reliable neural correlates of the psychedelic experience in humans consist of broadband reductions of spontaneous brain oscillatory activity (especially in the alpha range) and increased signal diversity and complexity, measured with electroencephalography (EEG) and magnetoencephalography (MEG) (Riba et al., 2002; Muthukumaraswamy et al., 2013; Kometer et al., 2013; Schenberg et al., 2015; Schartner et al., 2017; Valle et al., 2016; Carhart-Harris et al., 2016; Timmermann et al., 2019). When measured using functional magnetic resonance imaging (fMRI), psychedelic experiences are associated with global alterations in the functional connectivity between distinct functional networks (Carhart-Harris et al., 2012; Roseman et al., 2014; Tagliazucchi et al., 2016; Carhart-Harris et al., 2016; Müller et al., 2018; Preller et al., 2018). These changes could be attributed to the activation of layer 5 pyramidal neurons with high density of 5-HT_2A_ receptors, the main target site of psychedelics (Muthukumaraswamy et al., 2013; Nichols, 2016; Deco et al., 2018; Kringelbach et al., 2020). Activating these cells may impact spike-wave coherence, causing a disordering or ‘scrambling’ effect that manifests at the population and subjective levels as an increase in the entropy of spontaneous activity and as a relaxation of the high-level priors or beliefs (Carhart-Harris et al., 2014; Carhart-Harris, 2018; Carhart-Harris and Friston, 2019). In particular, this hypothesis is consistent with fMRI experiments showing that psychedelics decrease the intrinsic functional connectivity of the default mode network (DMN), a set of regions involved with self-referential processing and rich with 5-HT_2A_ receptors (Carhart-Harris et al., 2012; Palhano-Fontes et al., 2015; Preller et al., 2018). A consistent effect is seen in relation to other high-level intrinsic brain networks (Carhart-Harris et al., 2016).

However, all of the above listed studies were conducted in controlled laboratory environments, restricting the range of contextual or non-pharmacological variables surrounding the experience itself, and limiting the ecological validity of these experiments and their findings (Shamay-Tsoory et al., 2019). Since psychedelic experiences are very sensitive to the pre-existent individual state of mind (“set”) and surrounding environment (“setting”), it is difficult to estimate to which extent knowledge gathered in hospitals and research facilities can be readily generalized to natural or ecological settings (Hartogsohn, 2016; Carhart-Harris et al., 2018). As shown in a meta-analysis of 409 psilocybin administrations to 261 healthy subjects, anxious reactions are strongly associated to experiments conducted in neuroimaging facilities (Studerus et al., 2012), in agreement with the recommendation to avoid overly clinical environments in human psychedelic research (Johnson et al., 2008). Moreover, a large prospective survey (Haijen et al., 2018) showed that having clear individual motivation for the experience (“intention”) predicted the occurrence of mystical-type or peak experiences (Griffiths et al., 2006), with potential long-term positive effects in well-being after the experience (Griffiths et al., 2008); also, feeling well prepared and comfortable in the chosen setting was associated with less challenging experiences, as well as with higher well-being scores after the experience. It has been argued that potentially harmful effects could arise when experiments are conducted in unfavorable contexts, and that some early studies ignored or even manipulated these non-pharmacological factors with the purpose of eliciting unpleasant experiences (Carhart-Harris et al., 2018).

Taking heed of these observations, modern research with psychedelics is attentive to set and setting, which could represent an important source of success over past endeavors. It is recommended that subjects receive extensive psychological preparation, and that experiments be conducted in aesthetically pleasant, well-regulated settings (Johnson et al., 2008; Carhart-Harris et al., 2018; Kaelen et al., 2018). In spite of these precautions, research conducted in controlled laboratory settings remains unnatural for most participants. Real-world psychedelic experiences often take place in nature, private houses, or retreats, with the latter featuring such influences as scents, music and chants derived from indigenous traditions. Moreover, many psychedelic experiences are, in fact, social experiences undertaken by a group of interacting individuals who are supervised by a facilitator, guide or ‘shaman’(Carhart-Harris et al., 2018; Johnson et al., 2019). Additionally, real-world psychedelic use is often planned in advance, reflecting pre-existing motivations that may be difficult to replicate in double-blind placebo-controlled laboratory experiments (Haijen et al., 2018). Studies conducted in natural settings, i.e. settings that have been chosen by subjects beforehand and without interference from the researchers, are beginning to emerge as an alternative way to investigate the psychedelic experience *in situ*. Examples include studies of naturalistic ayahuasca (Kuypers et al., 2016), psilocybin (Prochazkova et al., 2018), and 5-MeO-DMT use (Uthaug et al., 2019).

Experiments conducted in natural settings could also contribute to the increasing body of knowledge on the neural correlates of the psychedelic experience, considering the substantially different conditions of laboratory experiments and their potential impact on the state of mind of the participants. To date, this line of research has received little to none attention. While some studies have attempted brain activity recordings in natural settings, they have been severely limited by underdeveloped mobile EEG technology, low sample sizes, and the challenge of collecting clear signal in such contexts (Don et al., 1998; Stuckey et al., 2005; Acosta-Urquidi, 2015). However, this technology has advanced considerably in recent years, allowing research-grade EEG recordings with fully wireless and minimally invasive headsets. Thus, this approach now appears to be ripe for field recordings of psychedelic use (Marini et al., 2019).

We recorded neural activity and subjective effects from 35 subjects who inhaled DMT in freebase form in their preferred context of DMT use (details below). Our experiment was designed to obtain good quality EEG recordings while simultaneously minimizing external disruptions in the subjects’ experiences. We chose to investigate experiences induced by DMT due to its intense but short-lasting effects (which drastically reduced the total duration of the experiment) (Szara, 1956; Strassman et al., 1994; Shulgin and Shulgin, 1997), and also since only one report to date investigated the effects of DMT alone (i.e. without combination with monoamine oxidase inhibitor [MAOIs]) on neural activity recordings acquired during controlled laboratory conditions (Timmermann et al., 2019). DMT is frequently consumed for exploratory and ceremonial or spiritual reasons, either alone in its freebase form, or crystallized over non-psychoactive plant leaves and then inhaled after combustion, which represents a comparatively understudied route of administration (Cakic et al., 2010; Winstock et al., 2014). We assessed the subjective effects directly after each experience, including factors related to the typical effects of psychedelic drugs (including mystical-type experience) as well as non-pharmacological factors related to environmental variables and social group interactions.

## Materials and methods

### Participants

Thirty-five participants (7 females; 33.1 ± 6 years; 92.2 ± 201.4 previous experiences with ayahuasca; 3.6 ± 5.6 previous experiences with DMT alone) were recruited by word-of-mouth and social media advertising between May and December 2019. All participants were asked to call to a number provided in the experiment advertisement, and were instructed to remain anonymous if preferred. During this first conversation participants were briefed on the details and objective of the experiment, as well as on the inclusion and exclusion criteria, and were offered an electronic version of the informed consent form. After checking the criteria to participate in the study (to be detailed below), subjects informed the location, date and time chosen for their DMT use. Four members of the research team, including a psychiatrist and a clinical psychologist, attended the appointment and presented the subject with an informed consent form. After signature, subjects underwent a psychiatric interview to screen for the exclusion criteria that are detailed in the following section.

This study was conducted in accordance with the Helsinki declaration and approved by the Committee for Research Ethics at the ‘Jose Maria Ramos Mejia’ General Hospital (Buenos Aires, Argentina).

All research data associated with this manuscript is publicly available (10.5281/zenodo.3992359)

### Inclusion and exclusion criteria

Participants were required to have at least two previous experiences with ayahuasca or DMT, abstain from consuming psychoactive drugs (including alcohol, caffeine and tobacco) at least 24 hours prior to the study, and to be willing to engage in their preferred use of DMT in the presence of four research team members. Researchers did not provide DMT or other psychoactive compounds to the subjects, nor interacted with their use of the substance.

All subjects were aged between 21 and 65 years, and pregnant women were excluded from the experiment. Subjects who declared past difficult experiences with psychedelics with lasting negative psychological sequelae or who put themselves or others at risk were excluded from the experiment. A non-diagnostic psychiatric interview (SCID-CT; First, 2014) was conducted according to the guidelines by Johnson et al. (2008). Subjects who fulfilled DSM-IV criteria for the following disorders were excluded from the experiment: schizophrenia or other psychotic disorders, and type 1 or 2 bipolar disorder (both also in first and second degree relatives), substance abuse or dependence over the last 5 years (excluding nicotine), depressive disorders, recurrent depressive episodes, obsessive-compulsive disorder, generalized anxiety disorder, dysthymia, panic disorder, bulimia or anorexia, as well as subjects with history of neurological disorders. Subjects who presented one standard deviation above the mean in the State-Trait Anxiety Inventory (Spielberger, 2010) were excluded, as well as subjects under psychiatric medication of any kind.

### Setting and DMT administration

Before and after DMT administration, all participants completed a series of questionnaires to be detailed in the following section.

Subjects consumed DMT in their preferred or usual setting, including their own choice of music, scents, lightning, and other contextual factors. After being fitted with the EEG cap, the subjects were briefed on the procedure for baseline recordings (see next session), and were instructed to keep their eyes closed, relax and maintain an upright sitting position, avoiding head movement and jaw clenching to prevent muscle artifacts. Subjects were asked to wait approximately five minutes after feeling a return to baseline to indicate the end of the recording, even though they could do so before if desired.

Only four subjects self-administered DMT, all others were assisted by a facilitator. After receiving instructions from their facilitators and, in some cases, practicing breath exercises, subjects gradually inhaled the smokes and vapors resulting from the combustion of freebase DMT, in all cases recrystallized over non-psychoactive plant leaves; the leaves employed by the participants for this purpose comprised white sage (*Salvia apiana*) and common jasmine (*Jasminum officinale*). Facilitators withdrew the pipe when subjects either stopped inhaling and leaned back, or exhausted the contents of the pipe.

The average load of the pipes was estimated by the participants or their facilitators at 40 mg freebase DMT. In all cases DMT was extracted from the root of *Mimosa hostilis* (also known as *Mimosa tenuiflora* or jurema) (Ott, 1999). The presence of DMT was verified in all samples by high performance liquid chromatography coupled to mass spectrometry for profiling and qualitative analysis.

### Assessment of set, setting and psychedelic experience

Participants completed a series of questionnaires immediately before and after their DMT experience. Before the DMT experience, participants completed Spanish versions of the State-Trait Anxiety Inventory (STAI trait) (Spielberger, 2010), and questions introduced in Table 3 of Haijen et al. (2018) to assess the self-reported adequateness of set, setting and intentions. After the DMT experience, participants completed the 5D altered states of consciousness scale (5D-ASC) (Studerus et al, 2010), the mystical experience questionnaire (MEQ-30) (Barrett et al., 2015), the near-death experience scale (NDE) (Greyson, 1983), and a series of questions to assess the impact of set, setting and social interactions on the psychedelic experience (post-experience questionnaire, or “Post”). Immediately before and after the DMT experience, participants completed Spanish versions of the Big Five personality (BFI) test (John and Srivastava, 1999), STAI state (Spielberger, 2010), and Tellegen absorption scale (TAS) (Tellegen and Atkinson, 1974). These questionnaires are described with more detail below.

#### STAI trait and state

Commonly used measure of trait and state anxiety, each comprising 20 items.

#### Set, setting and intentions

12 items assessing non-pharmacological contextual factors prior to psychedelic experiences. Using principal component analysis, Haijen et al. showed that items can be clustered into three components using the loadings presented in Table 3 of Haijen et al. (2018): ‘set’, ‘setting’, and ‘clear intentions’.

#### 5D-ASC

94 items assessing different aspects of altered states of consciousness, understood as temporary deviations from normal waking consciousness. Consists of five dimensions, each with multiple lower-order scales. The dimensions and their corresponding scales are as follows: oceanic boundlessness (experience of unity, spiritual experience, blissful state, insightfulness), anxious ego dissolution (disembodiment, impaired control and cognition, anxiety), visionary restructuralization (complex imagery, elementary imagery, audio-visual synesthesia, changed meaning of percepts).

#### MEQ-30

30 items from which four subscale scores are computed: mystical, positive mood, transcendence of time and space, and ineffability, which are considered the most relevant and defining aspects of mystical-type experiences. We extended these four subscales with ‘awe’, a construct derived from MEQ-30, characterizing a profound emotion in the presence of a vast or overwhelming stimulus that requires accommodation of mental structures, and proposed as a mediator of the downstream therapeutic effects of psychedelics (Hendricks et al., 2018).

#### NDE

16 items assessing alterations in consciousness arising from near-death experiences. Divided into four components: cognitive, affective, paranormal and transcendental. Its use was motivated by multiple previous reports likening the acute effects of DMT to near-death experiences (Strassman, 2000; Timmermann et al., 2018; Martial et al., 2019).

#### Post

27 items assessing the influence of non-pharmacological contextual factors after the experience, divided into four groups: set, setting, social and fusion.

#### BFI

44-item Big Five inventory, measuring participants on five dimensions of personality (extraversion, agreeableness, conscientiousness, neuroticism, openness).

#### TAS

34-item scale developed to measure the capacity of an individual to become absorbed in the performance of a task.

### EEG acquisition

EEG data were recorded with a 24-channel mobile system (mBrainTrain LLC, Belgrade, Serbia; http://www.mbraintrain.com/) attached to an elastic electrode cap (EASYCAP GmbH, Inning, Germany; www.easycap.de). Twenty-four Ag/AgCl electrodes were positioned at standard 10– 20 locations (Fp1, Fp2, Fz, F7, F8, FC1, FC2, Cz, C3, C4, T7, T8, CPz, CP1, CP2, CP5, CP6, TP9, TP10, Pz, P3, P4, O1, and O2). Reference and ground electrodes were placed at FCz and AFz sites. The wireless EEG DC amplifier (weight = 60 g; size = 82 × 51 × 12 mm; resolution = 24 bit; sampling rate = 500 Hz, 0–250 Hz pass-band) was attached to the back of the electrode cap (between electrodes O1 and O2) and sent digitized EEG data via Bluetooth to a Notebook held by a experimenter sitting behind the participant.

Prior to the administration of DMT, baseline EEG activity was acquired with eyes open and closed (five minutes each). After the DMT was administered, EEG recordings started when subjects exhaled, and lasted until the subject indicated a return to baseline (6 ± 1.4 min).

### EEG preprocessing and analysis

EEG data was preprocessed using EEGLAB (https://sccn.ucsd.edu/eeglab/index.php) (Delorme and Makeig, 2004) with the following procedure. Data was divided into 2 s. epochs and the first 5 epochs were removed for all subjects and conditions. Data was bandpass-filtered (1 – 90 Hz) and notch-filtered (47.5 – 52.5 Hz). Artifact-laden channels were first detected using EEGLAB automated methods, which reject channels computing kurtosis (threshold = 5), probability (threshold = 5) and the rejection of channels with ±2.5 standard deviations from the mean in any parameter (4.4 ± 1.8 channels removed per subject). All channels were manually inspected before rejection (mean 30% rejected channels, max. 8 channels) and then interpolated using data from the surrounding channels. Epochs to be rejected were flagged automatically and in all cases removed after manual inspection (21.3 ± 13.7 epochs rejected per subject). Infomax independent component analysis (ICA) was then applied to data from each individual participant, and used to manually identify and remove components related eye movements, blinking and muscle artifacts based following a semi-automatic procedure based on the frequency content and scalp topography of each component (Brunner et al., 2013), which was subsequently corroborated by visual inspection (2.7 ± 1.1 components removed). According to the previously defined criteria, 6 subjects were discarded from the subsequent EEG analysis due to an excessive number of rejected epochs and/or channels, resulting in 29 subjects for subsequent analysis.

The logarithmic power spectral density (LPSD) in the delta (1–4 Hz), theta (4–8 Hz), alpha (8– 12 Hz), beta (12–30 Hz) and gamma (30–40 Hz) bands was computed for each subject, condition and channel using a fast Fourier transform with a Hanning-tapered window, as implemented in EEGLAB. The time-frequency decomposition was obtained using the “newtimef.m”EEGLAB function. 2 s epochs were convolved using Morlet wavelets to yield time-frequency spectrograms for each subject and condition, with 80 frequency bins (< 40 Hz).

### Coherence and metastability

For each EEG channel, a 4th order Butterworth bandpass filter was applied to obtain the narrow band signals in the alpha to gamma bands. Afterwards, a Hilbert transform was applied to obtain the analytical representation of the filtered time series. Given a real-valued signal x(t), the Hilbert transform yields a complex representation given by z (t) = u(t) + iv (t) = x (t) + iH (x (t)). From this representation, the instantaneous amplitude can be computed as 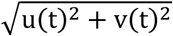, and the instantaneous phase as 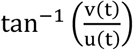.

The instantaneous amount of synchrony between bandpass-filtered channels can be obtained from the Kuramoto order parameter as 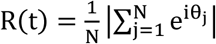, where the sum is over signal samples and θ_j_ represents the instantaneous phase at the j-th sample. R(t) ranges between 0 and 1, indicating minimum and maximum synchrony, respectively (Cavanna et al., 2018; Vivot et al., 2020). The metastability (K) is computed as the variance of R(t), and represents the amplitude of the dynamical repertoire of the channels (i.e. the occurrence of transient synchronization/de-synchronization between the signals) (Shanahan, 2010). The coherence (C) is computed as the mean of R(t), and indicates the average level of synchronization between channels.

### Lempel-Ziv complexity

Following previous work assessing signal diversity by means of algorithmic complexity (Schartner, et al., 2017; Timmermann et al., 2019), we estimated broadband signal complexity using the Lempel-Ziv lossless compression algorithm to binary time series obtained from a median split of the instantaneous amplitude (obtained via Hilbert transform) after Z-score conversion. We avoided biases due to unbalanced sequences since by definition of the median split all channels presented the same proportion of 1’s and 0’s. The Lempel-Ziv algorithm divides the binary string into non-overlapping and unique binary substrings. More substrings are required as a function of signal diversity. The total number of these substrings defines the Lempel-Ziv complexity, which is maximal for a completely random sequence and minimal for a completely regular sequence.

### Statistical analysis

Paired t-tests were used to compare psychometric (before vs. after) and EEG (DMT vs. eyes open/eyes closed) results, determining statistical significance either at p<0.05 after Benjamini-Hochberg false discovery rate (FDR), or at p<0.05 after Bonferroni correction for multiple comparisons (depending on the total number of comparisons, indicated in each particular case). Correlational analyses between EEG features and the scores of questionnaires were performed using Pearson’s linear correlation coefficient, and reported significant at p<0.05 after FDR correction.

## Results

### Set, setting and psychedelic experience

We first measured the self-reported adequateness of set, setting and intentions applying the item loadings presented in Haijen et al. (2018), resulting in 71.43% ± 27.36% for ‘set’, 88.24% ± 10.08% for ‘setting’ and 53.87% ± 23.78% for ‘clear intentions’, reported as mean % (over maximum score) ± SD.

Figure 1a summarizes the results from the scales used to assess the subjective effects (5D-ASC, MEQ-30, Post, and NDE). The same information is also provided in Table 1, where the mean % (over maximum score) ± SD are shown for all scales. The highest scores of the 5D-ASC questionnaire corresponded to three items in the visionary restructuralization dimension (elementary imagery, complex imagery, and audio-visual synesthesia) and to the blissful state item of the oceanic boundlessness dimension. Conversely, subjects scored low values in the impaired control and cognition, and anxiety items of the anxious ego dissolution dimension. The NDE scale presented the highest score in the affective component.

**Table 1.** Mean % (over maximum score) ± SD for all scales (5D-ASC, NDE, MEQ-30, Post) assessed immediately after the DMT experience.

| Scale | Factor | mean (%) | sd (%) |
| --- | --- | --- | --- |
| 5D-ASC | Unity | 47.53 | 29 |
|  | Spiritual | 49.61 | 13.92 |
|  | Blissful | 61.77 | 25.9 |
|  | Insightfulness | 45 | 22.32 |
|  | Disembodiment | 47.58 | 31.3 |
|  | Impaired | 22.21 | 18.98 |
|  | Anxiety | 12.06 | 19.5 |
|  | Complex imagery | 50.21 | 20.4 |
|  | Elementary imagery | 85.27 | 20.72 |
|  | Audiovisual | 56.32 | 28.35 |
|  | Changed meaning | 26.62 | 24.01 |
| NDE | Cognition | 26.17 | 24.32 |
|  | Affect | 60.94 | 26.39 |
|  | Paranormal | 32.03 | 21.64 |
|  | Transcendence | 31.64 | 25.72 |
| MEQ-30 | Mystical | 3.34 | 1.88 |
|  | Positive mood | 10.05 | 2.75 |
|  | Transcendence | 11.05 | 4.24 |
|  | Ineffability | 24.63 | 7.74 |
|  | Awe | 46.29 | 14.57 |
| POST | Setting | 76.48 | 25.38 |
|  | Social | 61.79 | 3.91 |
|  | Fusion | 35.93 | 9.21 |

**Figure 1.**
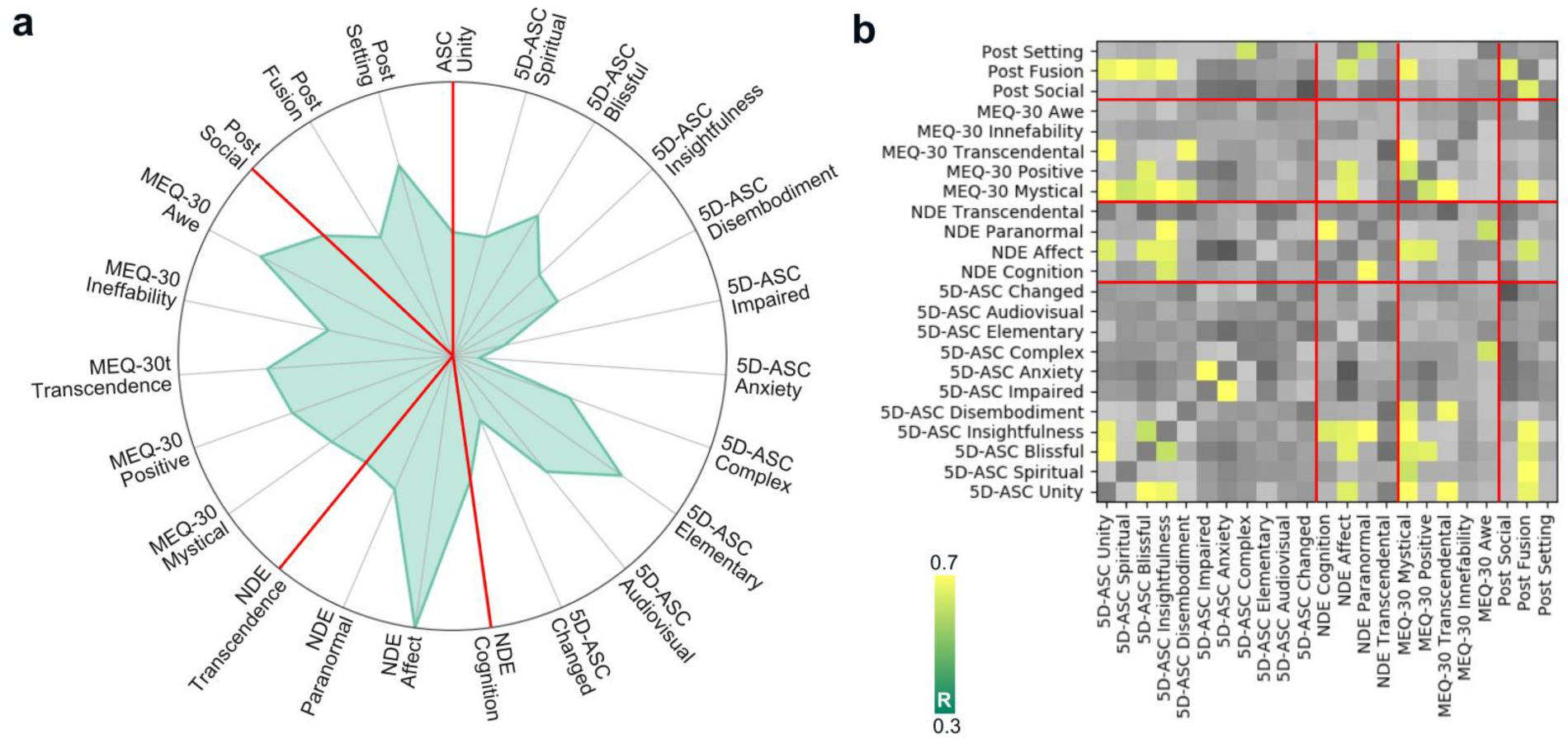
Subjective effects and their correlations. (**a)** Radar plot showing the normalized score of the 5D-ASC, MEQ-30, NDE and Post items, separated by red lines. Red lines divide the results from different questionnaires. (**b)** Matrix of Pearson’s linear correlation coefficients between all items of the three questionnaires, thresholded at p<0.05 (FDR-corrected) and R>0.3.

Figure 1b shows significant positive correlations between 5D-ASC items corresponding to the ‘oceanic boundlessness’ dimension and items from the other two scales that are related to the occurrence of mystical-type experiences: the affective component of the NDE scale (related to positive mood) and the ‘mystical’, ‘positive mood’, ‘transcendence of time and space’ (MEQ-30), and the ‘Fusion’ item of the Post. We also observed a significant positive correlation between the affective component of the NDE scale and the ‘positive mood’ and ‘mystical’ items of MEQ-30 and ‘Fusion’ (Post). Within-scale correlations also occurred for those items associated with mystical-type experiences: ‘experience of unity’, ‘spiritual experience’, ‘blissful state’ and ‘insightfulness’ for 5D-ASC; and ‘mystical’, ‘positive mood’, and ‘transcendence of time and space’ (MEQ-30). We also observed positive correlations between the relevance of social setting ‘fusion’ items (Post), and between anxiety and impaired control and cognition of the 5D-ASC.

We followed a criterion previously introduced to determine complete mystical-type experience, given by a score of 60% of the total possible score on each dimension of the MEQ-30 questionnaire (Barrett and Griffiths, 2017). According to this criterion, 13 of the 35 (37%) participants presented a complete mystical-type experience.

Next, we assessed the questionnaires that were obtained before and after the DMT experience (Table 2), which included personality assessment (BFI), anxiety (STAI state) and absorption (TAS), finding significant increases in ‘agreeableness’ (BFI) and absorption (TAS), and significant decreases in anxiety (STAI state).

**Table 2.** Comparison pre vs. post DMT for BFI, STAI and TAS (*p<0.05, Bonferroni-corrected).

| Scale | Factor | Pre |  | Post |  | t-value | p-value |
| --- | --- | --- | --- | --- | --- | --- | --- |
|  |  | mean | sd | mean | sd |  |  |
| BFI | Extraversion | 28.3 | 4.3 | 29.3 | 4.4 | -2.224 | 0.033 |
|  | Agreeableness | 34.5 | 4.6 | 35.7 | 4.1 | -3.072 | 0.004* |
|  | Conscientiousness | 28.8 | 6.1 | 29.3 | 6.2 | -1.382 | 0.176 |
|  | Neuroticism | 19.9 | 5.2 | 19.4 | 4.9 | 0.867 | 0.392 |
|  | Openness | 42.9 | 3.5 | 43.1 | 4 | -0.468 | 0.643 |
| STAI | State | 11.2 | 8 | 7.37 | 5.9 | 3.362 | 0.002* |
| TAS | Absorption | 16.1 | 6.6 | 18.2 | 8 | -3.788 | 0.001* |

Finally, we tested whether baseline questionnaire scores influenced the reported experience. Following Haijen et al. (2018), we found a significant positive correlation between the ‘clear intentions’ score and the ‘mystical’ MEQ-30 factor (R=0.36, p=0.029); we also found that baseline trait absorption correlated with the ‘mystical’ MEQ-30 factor (R=0.5, p=0.005), and that neuroticism (BFI) was predictive of ‘impaired control and cognition’ (R=0.44, p=0.007), but not of acute ‘anxiety’ (R=0.22, p=0.19) (5D-ASC). Baseline state anxiety also predicted the 5D-ASC items associated with challenging experience: ‘impaired control and cognition’ (R=0.41, p=0.012) and ‘anxiety’(R=0.65, p=0.001).

### EEG spectral power

We compared the LPSD averaged across all channels between the eyes open, eyes closed and DMT conditions; results are shown in Fig. 2a. The alpha peak at 10 Hz was significantly attenuated (p<0.05, FDR-corrected) for the DMT condition and, as expected, for the eyes open condition relative to the eyes closed condition. DMT also significantly increased (p<0.05, FDR-corrected) power corresponding to low (< 3 Hz) and high (> 36 Hz) frequencies. No significant differences were found between the DMT and eyes open condition.

**Figure 2.**
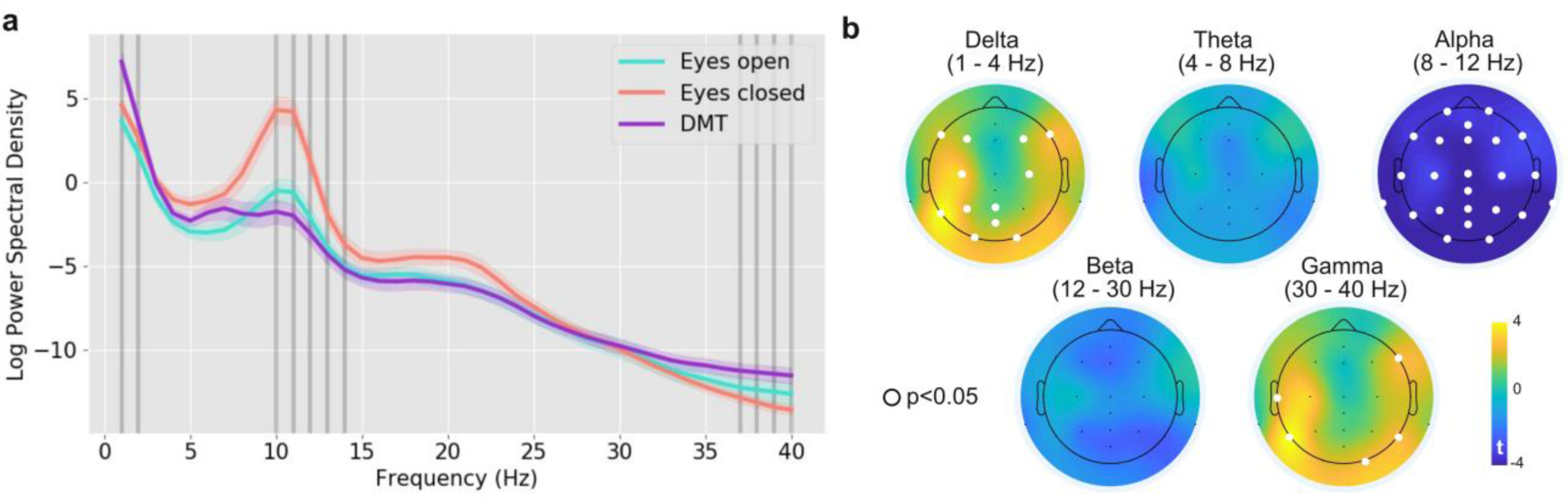
Frequency-specific changes in spectral power induced by DMT. **(a)** LPSD in 1 Hz bins for the eyes open, eyes closed and DMT conditions (mean ± SEM). Vertical bars (dark grey) indicate bins with significant differences between DMT and eyes closed (p<0.05, FDR-corrected). **(b)** Topographical maps for the t-values resulting from comparing DMT vs. eyes closed per frequency band. Electrodes in white represent significant differences with p<0.05 (FDR-corrected).

Power decreases under DMT relative to eyes closed were most widespread for the alpha band, for which all electrodes presented significant decreases (Fig. 2b). Consistently with the LPSD plot in Fig. 2a, we also observed more distributed power increases in the delta and gamma bands, restricted to electrodes located primarily in occipital, parietal and anterior-central regions (delta band), and in occipital, parietal and temporal regions (gamma band). We did not observe significant changes in EEG power for the theta and beta bands.

For each frequency band, we correlated the absolute and relative (i.e. minus eyes-closed) EEG power with the number of removed channels, epochs and ICA components, without observing significant correlations between both variables.

### Time-frequency analysis

We then compared the temporal evolution of spectral power during the first 7 minutes after DMT inhalation. We did not extend this analysis beyond this limit since several subjects started to indicate their return to baseline after this point. Fig. 3a presents the average time-dependent LPSD for all participants minus the baseline (eyes closed), and shows the progressive return to baseline topographic maps for the alpha band. These maps present average LPSD changes between 8 and 12 Hz at 40 s intervals, starting from the first 40 s after DMT administration (when the predominant blue indicates widespread reductions in LPSD compared to the eyes closed condition) and ending at ≈ 7 min (when the predominant red indicates that LPSD differences vs. eyes closed were close to zero in all channels).

**Figure 3.**
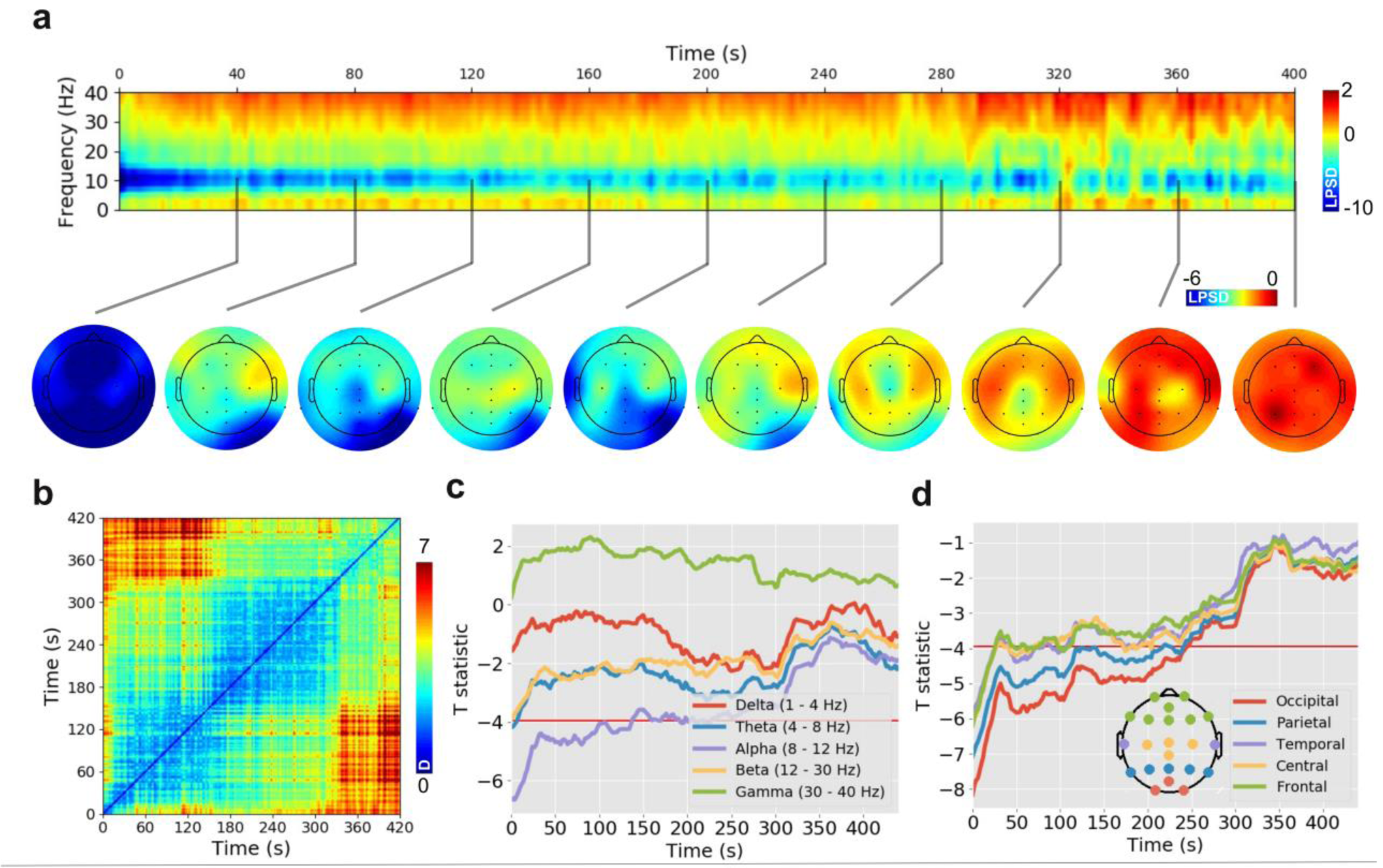
Time-resolved changes in EEG power reveal the gradual return to baseline after DMT administration (t=0). **(a)** Average LPSD for all participants in the DMT condition with baseline (eyes closed) subtraction. The scalp topographies show average LPSD changes in the alpha band (8-12 Hz) every 40 s. **(b)** Distance matrix for alpha power scalp topographies computed at different times after DMT administration. Each row and column indicates a time, and the matrix element contains the Euclidean distance (D) between the average alpha power for all electrodes at those two times. **(c)** T-value for the comparison DMT vs. baseline (eyes closed) for all frequency bands as a function of time after the DMT administration. The red horizontal bar indicates the threshold for statistical significance (p<0.05, FDR-corrected). **(d)** T-values for alpha power (DMT vs. eyes closed) vs. time, averaged at electrodes corresponding to different scalp locations.

The temporal evolution of alpha power is also manifest in the distance matrix for alpha power scalp topographies computed at different times after DMT administration (Fig. 3b), in analogy with the temporal generalization method proposed by King and Dehaene (King and Dehaene, 2014). In this matrix, each row and column indicates a time, and the matrix element contains the Euclidean distance between the average alpha power for all electrodes at those two times. Starting from t=0 (first column) the matrix elements turn progressively to red, indicating the gradual divergence of the scalp topography. In general, entries close to the diagonal presented the lowest distance values, while off-diagonal blocks presented the opposite behavior. This matrix structure is indicative of a smooth temporal evolution of average alpha power until 7 min after DMT administration.

Figure 3c presents the t-value for the comparison DMT vs. baseline (eyes closed) for all frequency bands as a function of time after the DMT administration. Alpha power presented significant decreases (indicated with a red horizontal line) until ≈ 3 min after onset of DMT effects, and the associated t-values started to stabilize around zero at 7 min. Time-dependent significant differences were not observed for the other frequency bands.

As shown in Fig. 3d, alpha power did not present a homogeneous return to baseline across the scalp, but showed more persistent effects for electrodes in occipital and parietal regions. This is consistent with the visualization of average LPSD at different times in Fig. 3a, where blue colors (indicative of LPSD decreases) persisted for longer durations at occipital and parietal regions.

### Correlations between neural and subjective effects

We correlated the scores obtained from all questionnaires (5D-ASC, MEQ-30, NDE, Post) with the LPSD averaged across frontal (Fp1, Fp2, Fz, F7, F8, FC1, FC2), central (Cz, C3, C4), occipital (O1 and O2), temporal (T7, T8) and parietal channels (CPz, CP1, CP2, CP5, CP6, TP9, TP10, Pz, P3, P4). The results from this analysis are shown in Fig. 4a. We only found significant correlations between subjective effects spectral power for the beta and gamma bands. In the first case, the ‘anxiety’ item of 5D-ASC correlated positively with central beta power, and the ‘cognition’ component of the NDE scale correlated positively with occipital, central and frontal beta power. In the second case, the ‘experience of unity’, ‘disembodiment’ (5D-ASC), ‘cognition’ (NDE), ‘mystical’ and ‘transcendence of time and space’ (MEQ-30) correlated positively with occipital gamma power; ‘anxiety’, ‘complex imagery’ (5D-ASC) and ‘awe’ (MEQ-30) correlated positively with central gamma; ‘cognition’ (NDE) and ‘transcendence of time and space’ (MEQ-30) correlated positively with frontal gamma; and ‘cognition’ (NDE) correlated positively with temporal gamma.

**Figure 4.**
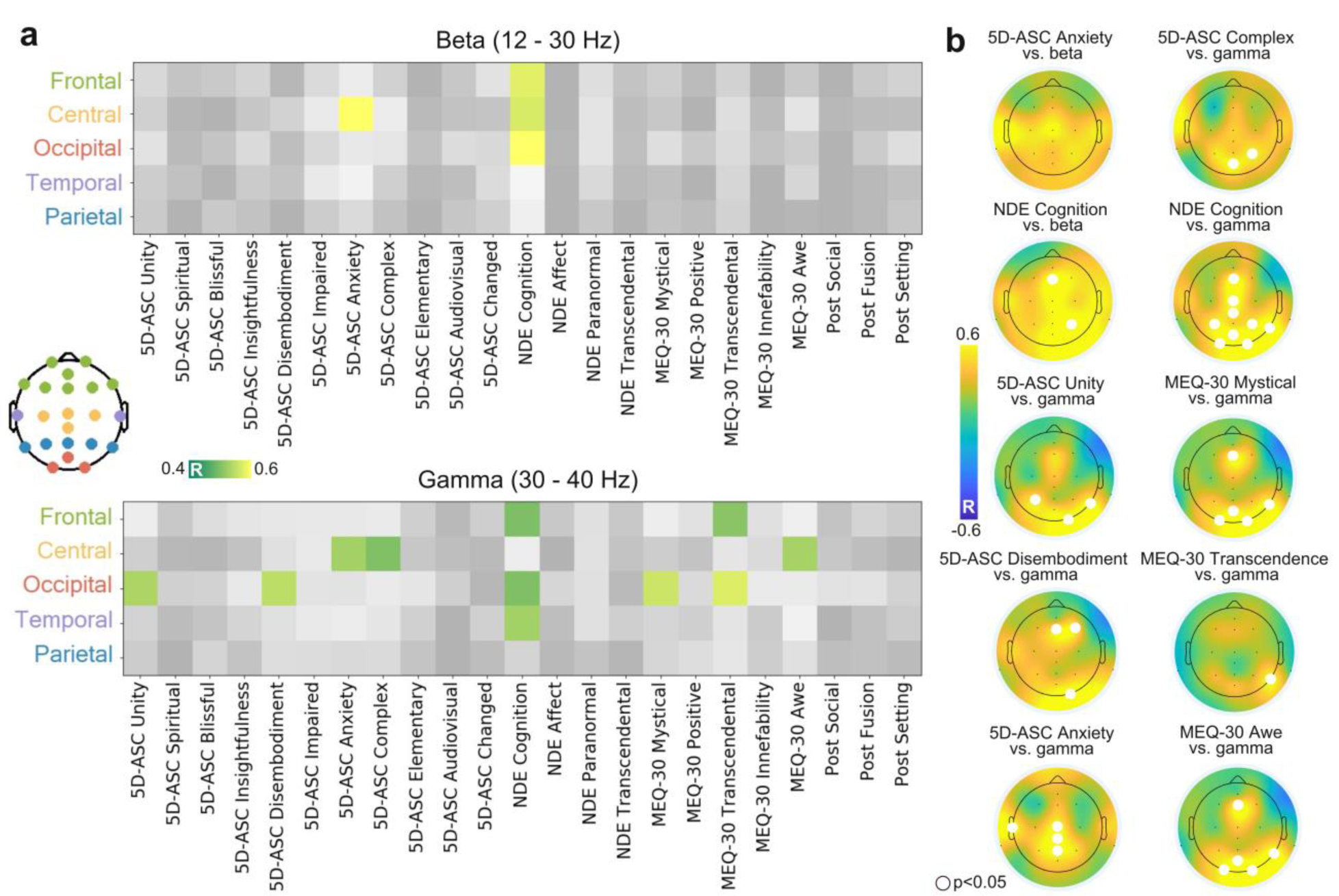
Correlations between neural and subjective effects. **(a)** Correlation between scores from all questionnaires with the LPSD averaged across frontal, central, occipital, temporal and parietal channels; significant correlations (p<0.05, FDR-corrected) were only found for the beta (up) and gamma (bottom) bands. **(b)** Scalp distributions of Pearson’s linear correlation coefficient between subjective effects and band-specific LPSD. Electrodes in white represent significant differences with p<0.05 (FDR-corrected).

Figure 4b shows scalp distributions of Pearson’s linear correlation coefficient between subjective effects and band-specific LPSD under the DMT condition. The power of beta oscillations presented significant correlations with ‘cognition’ (NDE), while the power of gamma oscillations presented significant correlations with ‘experience of unity’ (5D-ASC), ‘disembodiment’ (5D-ASC), ‘anxiety’ (5D-ASC), ‘complex visual imagery’ (5D-ASC), ‘cognition’ (NDE), ‘mystical’ (MEQ-30), ‘transcendence of time and space’ (MEQ-30), and ‘awe’ (MEQ-30). Significant channels were predominantly located in occipital and central regions.

We investigated whether the intensity of these correlations depended on time using the results from the time-frequency analysis (Fig. 5). We found variable temporal profiles of correlations between EEG power and reported subjective effects, with some variables presenting peak correlations at earlier times, e.g. ‘cognition’ (NDE) and beta/gamma power; ‘anxiety’ (5D-ASC) and gamma power, while other presented peak correlations at intermediate or later times, e.g. ‘experience of unity’, ‘disembodiment’, ‘impaired control and cognition’ (5D-ASC) and gamma power.

**Figure 5.**
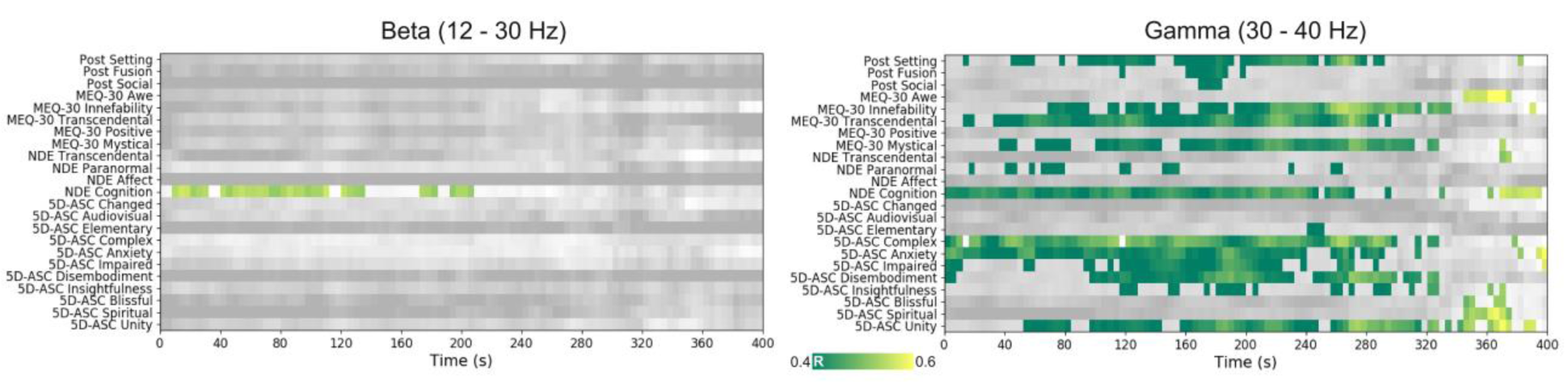
Correlation between subjective effects and time-resolved changes in beta and gamma power, thresholded at p<0.05 (FDR-corrected).

### Coherence, metastability and Lempel-Ziv complexity

Figure 6a presents the comparison of coherence (C) and metastability (K) between the DMT and eyes closed conditions. We found that after DMT administration both C and K decreased in the alpha band and increased in the gamma band. As shown in Fig. 6a, signal complexity (measured using the Lempel-Ziv algorithm) increased for all channels in the DMT condition vs. the eyes closed condition. These changes were similar in magnitude and distribution to those seen in the comparison of eyes open vs. eyes closed. We did not find significant correlations between the results of the psychometric questionnaires and EEG coherence, metastability or Lempel-Ziv complexity.

**Figure 6.**
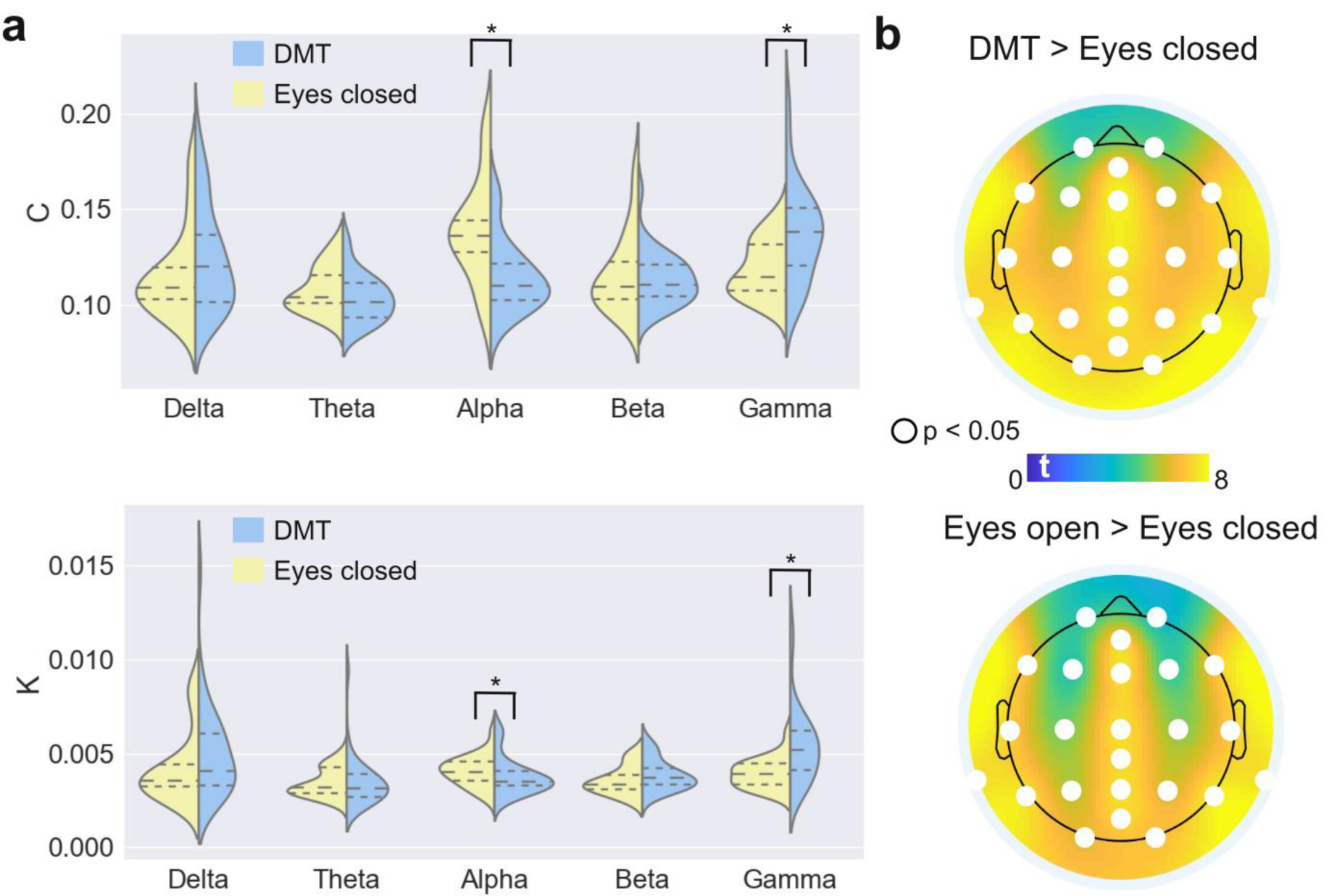
Changes in coherence (C), metastability (K) and signal complexity following DMT administration relative to the eyes closed baseline. **(a)** Violin plots for C (up) and K (bottom) computed from the DMT and eyes closed conditions. Long/short dashed lines represent the means and quartiles of the distributions (* p<0.05, FDR-corrected). **(b)** Signal complexity (Lempel-Ziv algorithm) t-value maps for DMT vs. eyes closed (up) and eyes open vs. eyes closed (bottom). Electrodes in white represent significant differences with p<0.05 (FDR-corrected).

## Discussion

We investigated the neural correlates of a DMT experience in a natural setting using modern wireless EEG technology. In particular, our study addressed the transient altered state of consciousness elicited by DMT alone, which to date has received less attention than the acute effects of psychedelics with longer half-lives (Barker, 2018). All participants had already decided to undergo an experience with DMT and were intrinsically motivated to do so; moreover, since they obtained the substance by their own means, we were able to recruit and study a comparatively large number of subjects - which is a particular merit of the present, naturalistic approach. Our results were largely consistent with previous studies conducted in laboratory settings, but also presented some differences which might hint of the potential effects of non-pharmacological contextual factors.

Conducting a search in the Altered States Database (http://www.asdb.info/) (Schmidt and Berkemeyer, 2018), we determined that the profile of subjective effects elicited by DMT in our experiment was comparable to previous reports. Changes in the ‘visionary restructuralization’ dimension were most prominent, followed by ‘oceanic boundlessness’ and ‘anxious ego dissolution’ (Gouzoulis-Mayfrank et al., 2005). Interestingly, we found high scores for the ‘affective’ component of the NDE questionnaire, in agreement with a recent related report (Timmermann et al., 2018), which is also a landmark feature of actual near-death experiences (Greyson et al., 1993). This result resonates with the proposal of endogenous DMT release when the brain undergoes severe stress (Strassman, 2000), even though this hypothesis has been severely criticized on the grounds of insufficient endogenous DMT concentration (Nichols, 2018). Concerning the MEQ-30 results, 37% of the participants fulfilled the criterion for a complete mystical experience, with motivation and trait absorption predicting the ‘mystical’ score, as previously reported in Haijen et al. (2018). This percentage is between 60% reported in multiple studies with psilocybin and 10% reported for LSD (Griffiths et al., 2006; Griffiths et al., 2008; Barrett and Griffiths, 2017). It must be noted, however, that the psilocybin studies were explicitly designed to favor mystical-type experiences, e.g. in samples of individuals interested in or studying comparative religion (Griffiths et al., 2006). A large online survey of ‘God encounter experiences’ found that a much larger proportion of experiences (73%) with DMT alone qualified as complete mystical; however, it is likely that due to the specific aim of the study the respondents had intentions of achieving this kind of experience (Griffiths et al., 2019). Given that psychedelics have previously been shown to enhance suggestibility (Carhart-Harris et al., 2015), and strong assumptions and increasing evidence for the role of expectation (Carhart-Harris et al., 2018) one must be mindful of the influence of priming on the reported content of psychedelic experiences.

Our study failed to replicate the long-standing association between openness and psychedelic experiences (MacLean et al., 2011; Lebedev et al., 2016; Bouso et al., 2018; Erritzoe et al., 2019). This association has been demonstrated for ayahuasca, psilocybin and LSD, even though multiple negative results have also been reported (Bouso et al., 2018). It must be emphasized that we conducted the corresponding questionnaires immediately after the DMT experience, which might reflect ‘state’ as opposed to ‘trait’ effects. Besides this point, we can speculate that the intense but brief experience induced by DMT could be insufficient to exert changes in this dimension of personality, even though future studies should assess this claim under more controlled experimental conditions. We also found an unexpected significant increase of the agreeableness dimension, which could be related to the social context of the experience, and hence overlooked by research conducted in different settings. Finally, subjects scored higher in absorption and lower in anxiety immediately after DMT compared with baseline, assessed prior to the experience.

Certain 5-HT_1A_ and 5-HT_1B_ receptor agonists reduce irritability (Olivier et al., 1994) and DMT (as most psychedelic tryptamines) activates these two serotonin receptor subtypes (Nichols, 2016; Ray, 2010; Zamberlan et al., 2018); however, reductions in anxiety observed following DMT administration, could relate to elevated preparatory levels of anxiety commonly observed prior to DMT experiences which are relieved once the experience has concluded and are thus are more related to anticipation than the actual effects of DMT. This highlights the need for a double-blind placebo-controlled paradigm to investigate potential anxiolytic effects of DMT. Increased absorption after psychedelic experiences in naturalistic settings has been previously demonstrated for ayahuasca (Bresnick and Levin, 2006) and for subthreshold doses of psychedelics (i.e. microdoses) (Polito and Stevenson, 2019). Our results confirm the close association between psychedelic effects and absorption, where the 5-HT_2A_ receptor has been specifically implicated (Ott et al., 2005).

While we replicated previous findings concerning decreased power in the alpha band (Riba et al., 2002; Muthukumaraswamy et al., 2013; Kometer et al., 2013; Schenberg et al., 2015; Schartner et al., 2017; Carhart-Harris et al., 2016; Timmermann et al., 2019), including work showing that this depends on 5-HT_2A_ receptor stimulation (Valle et al., 2016), some discrepancies appeared in the time course of alpha power changes after DMT inhalation in the present study. Reduced alpha power manifested most strongly immediately after DMT administration, and gradually receded towards baseline values after approximately 7 minutes. In contrast, previous work by Timmermann and colleagues reported that reports of subjective effect intensity were reduced to half of their peak value after 7 minutes and that alpha rhythm amplitude did not completely recover until approximately 15 minutes after the injection of doses ranging between 7 mg and 20 mg DMT fumarate (Timmermann et al., 2019). These differences in the time course of alpha power reductions could be attributed to different pharmacokinetics between inhaled and intravenous administration routes, with some reports suggesting an equally rapid onset but shorter duration in the case of inhaled DMT (Barker et al., 2018). Another possibility is that the effective dose in our study was lower than the doses employed by Timmermann and colleagues. Even though we did not quantify DMT content in the smoked samples, participants and facilitators declared a typical dose of 40 mg, in all cases extracted from *Mimosa hostilis* bark following a standard acid-base extraction procedure (Fasanello and Placke, 2007). Previous analysis of similar DMT samples showed an average purity of 85% (Riba et al., 2015), which would imply an average dose of 35 mg in our study, comparable to the maximum dose used in Timmermann, et al. (2019) when one considers brain availability from an inhaled dose vs. intravenous administration. While smoked and intravenous DMT are both more potent than intramuscular injections (Szara, 2007), more research is required to establish a more precise correspondence between these two routes of administrations. A third possibility is that subjects systematically underestimated the subjective effects at around 7 minutes compared with the peak of the experience, and erroneously declared a return to baseline in spite of still significant subjective and neurophysiological effects; however, the measured EEG power spectra appear to be incompatible with this possibility.

We observed more salient divergences with the previous report by Timmermann and colleagues at other frequency bands. The power of delta (1 – 4 Hz) oscillations increased compared with the eyes-closed baseline, while this increase only appeared as a trend in the report by Timmermann and colleagues. However, our results are consistent with an EEG study of the effects of ayahuasca, which found that delta power increases correlated with DMT plasma concentration, but did not correlate with the concentration of several psychoactive beta-carbolines also found in the brew (harmine, harmol, harmaline, harmalol and tetrahydroharmine) (Schenberg et al., 2015). Importantly, an independent EEG study of ayahuasca found small decreases in delta power that vanished at ≈ 120 min after ingestion, which coincides with the peak DMT plasma concentration (Riba et al., 2002). Psilocybin was found to increase lagged phase synchronization of delta oscillations (Kometer et al., 2015), and hypersynchronous delta activity has been reported in rodents after administration of DMT (Morley and Bradley, 1977) and 5-MeO-DMT (Riga et al., 2014), a close structural analogue of DMT (Shulgin and Shulgin, 1997); however, LSD decreased delta power in humans (Carhart-Harris et al., 2016). This heterogeneity could be in part explained by the different neurochemical profile of these drugs (Ray, 2010), especially by activity at other serotonin receptor subtypes (e.g. 5-HT_1A_, 5-HT_1B_) as suggested by 5-MeO-DMT-increased delta power in 5-HT_2A_ knockout mice (Riga et al., 2014). Also in contrast with the results reported by Timmermann and colleagues, we did not find that DMT increased power in the theta band, in agreement with other studies of ayahuasca, psilocybin and LSD. This discrepancy, however, could be caused by assessments of different contributions (oscillatory vs. fractal components) to the theta power (Timmermann et al., 2019).

Perhaps our most salient result consists of increased gamma power under DMT, correlating with multiple items from the 5D-ASC, NDE and MEQ-30 scales that reflect aspects of mystical-type experiences. Gamma power increases did not correlate with the number of discarded components/electrodes/channels, and were preserved after conservative EEG processing criteria which led us to discard 6 out of 35 participants. The role played by gamma oscillations in the neurobiology of psychedelic experiences remains unclear, mainly because gamma activity can result from muscle tension, jaw clenching and microsaccades (Fries et al., 2008). In contrast to the results reported by Timmermann et al. intravenous DMT, an early EEG field study of ayahuasca found hyper-coherence in the gamma band (Stuckey et al., 2005), consistent with the results from Schenberg and colleagues. However, we note that Timmermann et al. observed differences in the gamma band comparing baseline vs. DMT, but not when comparing placebo vs. DMT. In a study with 50 healthy participants, Kometer et al. found that psilocybin administration results in increased high frequency (55 – 100 Hz) oscillations within the high gamma range (Kometer et al., 2015) that correlated with the reported intensity of mystical experiences, also in line with a preliminary report that also found increased gamma global synchrony (Palenicek et al., 2016). MEG recordings acquired during the acute effects of LSD and psilocybin systematically failed to find gamma power increases compared with placebo (Muthukumaraswamy, et al. 2013, Carhart-Harris et al., 2016). Invasive recordings performed in animals support that psychedelic-induced gamma increases are at least partly mediated by 5-HT_2A_ activation (Goda et al., 2013). While past discussions on these discrepancies have focused on the possibility of EEG confounds within the gamma range, we believe that at least some of these discrepancies could also be attributed to differences in contextual factors. Variables related to set and setting could facilitate certain kinds of experiences whose neural correlates lie in the gamma range; moreover, anxious states may result in tension and movement, confounding activity in this band.

One of the most interesting characteristics of the psychedelic drugs is their capacity to induce peak experiences of unity and transcendence, combined with feelings of sacredness, deep meaning, positive mood, ineffability and paradoxicality (Griffiths et al., 2006; Griffiths et al., 2008; Barrett and Griffiths, 2017). These mystical-type experiences correlate with positive outcomes in therapeutic applications of psilocybin, occur with dose-dependent frequency, and are usually placed by research participants among the most meaningful experiences of their lives, even decades after the event (Griffiths et al., 2008; Griffiths et al., 2011; Garcia-Romeu et al., 2014; Griffiths et al., 2016). A possible neurophysiological mechanism underlying these experiences is increased local and long-range synchrony of gamma oscillations. Supporting this possibility, multiple studies have shown that states achieved by long-term meditators following certain traditions (e.g. non-dual awareness) (Lutz et al., 2004; Braboszcz et al., 2017; Berkovich-Ohana, 2017), share several defining features with the phenomenology of psychedelic-induced mystical-type experiences (Millière et al., 2018). It has also been proposed that surge of gamma band coherence near death is a correlate of NDE (Borjigin et al., 2013), an altered state of consciousness that has been compared with the DMT experience in terms of subjective experience (Strassman, 2000; Timmermann et al., 2018; Martial et al., 2019). In spite of their obvious relevance, very little is known about the neurobiological underpinnings of psychedelic-induced mystical-type experiences and is difficult to know to what extent contextual priming and framing plays a role in these experiences. Field studies seem especially apt to investigate the neurobiological basis of ‘mystical-type’, ‘peak’, or ‘unitive’ experiences, especially in the case of participants who approach psychedelic use from a spiritual perspective. We established gamma increases as a correlate of several 5D-ASC, NDE and MEQ-30 items related to the phenomenology of mystical experiences. As shown in Fig. 5, these correlations tended to manifest most strongly near the peak of the experience (≈ 3 min) and then receded gradually. Previous DMT research did not find correlations between gamma band oscillations and these items (Timmermann et al., 2018); however, this work assessed subjective experience using visual analogue scales (VAS) which might not be comparable with the results of well-validated psychometric questionnaires.

We replicated previous findings of increased signal diversity (as measured using the Lempel-Ziv algorithm) under the acute effects of serotonergic psychedelics, including DMT (Schartner et al., 2017; Timmermann et al., 2018). We also demonstrated that the collective properties of EEG oscillations differ under DMT in comparison to the baseline condition: alpha oscillations decreased in coherence and metastability, while gamma oscillations showed the opposite behaviour. As hypothesized previously by Stuckey and colleagues (Stuckey et al., 2005), and as supported by the discussion in the previous paragraph, gamma hyper-synchrony could be a manifestation of increased information binding underlying psychedelic-induced unitive experiences. Recently, we found increased low gamma band coherence and signal diversity in the EEG of expert meditators belonging to multiple different traditions (Vivot et al., 2020). These results are also consistent with a potential involvement of gamma oscillations in certain aspects of the psychedelic-experience that are both facilitated by natural settings and common to other non-pharmacological altered states of consciousness.

Perhaps one of the most important future roles of conducting studies in naturalistic settings is to understand how different contexts may contribute to the therapeutic properties of psychedelics. Again, this role is complementary to that of clinical trials designed to test the efficacy and safety of the treatment. It is becoming increasingly clear that non-pharmacological contextual factors cannot be easily disentangled from neurochemical action when it comes to the use of psychedelics in psychiatry (Carhart-Harris et al., 2018). In other words, the clinical merit of these drugs should be evaluated in relationship to the context where they are administered, and field studies could prove important to understand how non-clinical settings impact on psychological and neurophysiological variables that relate to the psychedelic experience and are of potential therapeutic value. It is also important to highlight that ‘underground’ use (i.e. outside of formal research settings) of psychedelic substances for therapeutic purposes is becoming increasingly popular, and that in those cases the experiences occur in settings very different to those seen in clinical facilities. Thus, conducting field studies is important to understand how psychedelics interact with the kind of environment where unregulated therapeutic uses are currently being pursued.

Our study presents multiple limitations inherent to observational field studies. The most obvious is the challenge of following a double-blind and placebo-controlled design, and thus our results could be influenced by expectation effects. Recent work has shown that a placebo could lead participants to think they underwent a psychedelic experience, even though the reported subjective effects differed substantially with those that are elicited by DMT (Olson et al., 2020). Also, the very strong and idiosyncratic effects of DMT render the possibility of similar acute effects as a consequence of placebo very unlikely. Investigating research participants with ample experience with DMT and/or ayahuasca (as in this study) could contribute to attenuate these effects. Another important limitation is lack of precise dose quantification. Even if quantitative assays could be performed in the DMT sample, combustion followed by inhalation is not completely efficient as a drug delivery method, resulting in an unknown total effective dose. As noted in Timmermann et al. (2019), however, exploring a range of different doses could facilitate the process of establishing correlations between neural activity data and psychometric questionnaires. Finally, due to privacy considerations we opted not to pursue follow-up measurements, thus prioritizing the anonymity of the participants and minimizing their contact with members of the research team. Because of these multiple limitations, we believe that field studies of psychedelic use should be considered as valuable alternatives in between massive online surveys and highly controlled laboratory experiments, and thus evaluated in light of their intrinsic merits, such as the possibility of investigating pre-planned psychedelic use in motivated and experienced individuals, and in the conditions where these substances are most frequently consumed, thereby raising the ecological validity of these findings.

In conclusion, we characterized the neural and subjective effects of DMT alone consumed in natural settings, reproducing known signatures of the psychedelic experience, and finding others that suggest interactions between large-scale rhythmic brain activity and non-pharmacological contextual factors. Our profile of the DMT experience complements both laboratory experiments and survey studies, and demonstrates the feasibility of obtaining neural activity recordings during psychedelic experiences in the field. Future studies should capitalize on the advantages of field studies to provide a more comprehensive and integral view of the psychedelic experience and its relation to neuronal activity, as well as to investigate the interaction between set, setting, intention and the therapeutic potential of psychedelics.

## Acknowledgments

We thank Gonzalo Sierra for his continuous support of this study. The authors also acknowledge Toyoko LLC for granting cloud computing services.

